# Eradicating Citrus Greening By Dual Action Targeting of *Liberibacter asiaticus* and *Diaphorina citri*

**DOI:** 10.1101/2022.02.07.479449

**Authors:** Catherine Farrell, Eric W. Triplett, Neil D. Theise, Cadance A. Lowell, Anthony R. Arment

## Abstract

Huanglongbing (citrus greening) is caused by the unculturable, gram negative bacterium *Liberibacter asiaticus* and transmitted by the Asian Citrus Psyllid (*Diaphorina citri*). Prior research demonstrated that the rifampicin-derivative TPR-1 as well as Palisades Therapeutic (PT) compounds PT159 and PT160 were effective at inhibiting growth of the *L. cresens*, the only culturable model for *Las*, showing 100% inhibition at 0.5 μg mL^−1^, and 80% inhibition at 0.05 μg mL^−1^.

Research with the PT glucocorticoid antagonist compounds demonstrated an inhibition of bacterial growth in *L crescens* as well, possibly a synergistic addition to TPR-1. A search of the *L. asiaticus* genome revealed the presence of 3 putative glucocorticoid response elements (GRE). Putative GRE are present in all organisms and may be involved in organism signaling as well as host-pathogen crosstalk.

In *Drosophila*, the estrogen-related receptor (ERR) transcription factor is a master regulator of larval and pupal maturation. ERR binds to GRE promoter elements in the genome to regulate the transcription of pathways involved in glucose metabolism. Null mutations of *ERR* fail to leave pupation and demonstrate elevated glucose levels when compared to wild-type. Treatment of *Drosophila* larvae with PT compounds demonstrated developmental delay disruptions in pupation, including premature pupation and increased lethality. Pupae that had been fed PT compounds as larvae showed elevated glucose levels when compared to controls. Sequences identified in *Drosophila ERR* showed high homology to 2 proteins in *D. citri* (ERR-like 1 and ERR-like 2), leading us to hypothesize that a glucocorticoid antagonist may disrupt psyllid development. Because these genes and signaling pathways are highly conserved between these taxa, we propose that *Drosophila* serves as a good model for screening potential compounds for the control of psyllid populations.

Collectively, we hypothesize that the dual action of these compounds offers a solution to HLB that can salvage the dying citrus industry in Florida and provide the first effective treatment. By simultaneous targeting of both *Las* and *D. citri*, it may be possible to both cure infected trees and block psyllid development in adults feeding on those same trees. Disruption of the psyllid’s ability to carry *Las* or cure *Las* from the psyllid gut are also both substantial possibilities.

## Introduction

Citrus is one of the top specialty crops in the United States (U.S.), making up a third of bearing area and accounting for 16% total value in U.S. fruit production (Li et al., 2020). Florida has traditionally been the largest orange and orange juice producer and has been most heavily impacted by Huanglongbing (HLB or citrus greening). HLB causes economic loss, because it affects both annual fruit quality and yield in addition to long term damage to the health of the tree. Trees infected with HLB typically can become unproductive in as few as 2-5 years, reducing an average 50-year lifespan to 7-10 years (Li et al., 2020). Estimated damages over the last 5-years are at $1 billion dollars per year, including a loss of 5000 jobs annually. Grove-bearing areas have declined by an estimated 30% since 2005 with an estimated 74% reduction in production (Court et al., 2017). HLB damage shifted the top citrus producer in 2016 from Florida to California. To date, no effective treatment or cure has been successfully employed.

HLB is caused by the unculturable, fastidious, gram negative, obligate bacteria *Liberibacter asiaticus* (Gottwald, 2010). *Liberibacter* spp. are phloem-limited and have been designated into three species based upon DNA and morphological analyses: *L. asiaticus (Las), L. africanus (Laf)* and *L. americanus (Lam)*. Transmission of these pathogens occurs through the feeding of parasitic psyllids. *Las* and *Lam* are transmitted by *Diaphorina citri* Kuwayama; Laf is transmitted by *Trioza erytrea* (Del Guercio) (Halbert, 2009). Progress has been made by Fagen et al. (2014) with the establishment of a closely related culturable species, *L. crescens*, from the sap of a mountain papaya in Puerto Rico. *L. crescens (Lc)* has 94.7% genetic identity of the 16S rRNA gene with both *Las and Lam*, making it a promising model for the study of *Liberibacter* pathology. As *Lc* is the only culturable relative, it a valuable tool for comparing the effectiveness of treatment regimens. Zhang et al. (2014) conducted a comparative screen of antibiotic effectiveness against *Las* using a graft-evaluation method; one of the most effective antibiotics tested was rifampicin. However, rifampicin is light-sensitive and not as thermally stabile as other antibiotics.

In *Drosophila*, a master developmental regulatory switch is the Estrogen-related Receptor protein (ERR), a type of glucocorticoid receptor. This nuclear receptor serves as a transcriptional regulator of glucose metabolism by binding to glucocorticoid response elements (GREs) upstream of several key genes necessary in the glucose metabolic pathway. Disruption of the ERR halts larval development and alters base metabolism to low ATP levels and elevated glucose (Kovalenko et al., 2019; Tennessen et al., 2011). The genome of *D. citri* revealed two genes (*ERR-like1* and *ERR-like2*) with high homology to *ERR* in *Drosophila*. It was hypothesized that this disruption could result in psyllid reduction/elimination by limitations placed on the lifecycle.

Collaborators at PT successfully synthesized a rifampicin derivative with enhanced thermal stability and light insensitivity. Other compounds from PT are well characterized glucocorticoid receptor antagonists. Our working model is that by pairing these compounds, a dual action synergy can be achieved that will both cure infected trees and disrupt the lifecycle of the *D. citri* transmission vector.

## Materials & Methods

### PT Compounds

Compounds PT150 and PT155 are glucocorticoid antagonists. TPR-1 is a modified rifampicin derivative. PT159 and PT160 are mixtures of PT150 and TPR-1 and PT155 and TPR-1. All compounds are proprietary under Palisades Therapeutics, LC. All compounds were dissolved and diluted in DMSO.

### Antibiotic Sensitivity in *L. crescens*

*Lc* was grown for three days to form an inoculant. Tested compounds were dispensed in quadruplicate into 96-well plates using an equal number of *Lc* and a media control; the total volume of each well was 200 μL. Plates were sealed with adhesive sheet covers and incubated with shaking for 3 days at 28°C at 150 rpm. After incubation, 90 μL of each well was mixed with 10 μL of Presto Blue (ThermoFisher Sci) in a clean, 96-well black plate. Plates were wrapped in foil at room temperature for 4 hours to develop. Plate wells were read by fluorescence to determine cell viability. Cell viability was calculated as the ratio of control to treatment absorbance values.

### Fruitfly Feeding

*Drosophila* used in this assay were selected during the third instar foraging stage. Larvae were transferred to grape juice agar plates, and fed compounds suspended in a solution of water with red food coloring. This solution was combined with dry yeast, and larvae were allowed to feed for 48 hours. At 48 hours, populations were assessed for survival and developmental stage.

### Preparation of Pupae for Glucose Assay

Pupae for glucose assay were ground in a microtube with a disposable pestle in 100μL of 0.01 MES, pH 6.0. Twenty microliters were used in each assay and performed in triplicate.

### Glucose Assays

The method of Huang et al. (2018) was adapted to a microplate assay format. Briefly, in each well, 10mM KMnO_4_ was reduced to MnO_2_ in 0.1M MES, pH6.0. Glucose standards along with larval samples were added to wells containing 0.321 mg mL^−1^ of glucose oxidase and incubated for 40 minutes at 37°C. Absorbance was measured at 346 to generate the standard curve and experimental data. Assays were performed in triplicate.

## Results

### *L. crescens* growth is inhibited by compounds PT159, PT160, and TPR-1

### BLAST Analysis

A protein blast analysis was conducted to compare the *Drosophila* ERR gene against the two genes in *D. citri* (ERR-like 1 and ERR-like 2). The expected values are described in the chart below.

**Table 1.** Homology Between Drosophila ERR and *D. citri* ERR-like Proteins

|  | mRNA | Protein | Expect Value |
| --- | --- | --- | --- |
| ERR-like 1 | XM_026831933 | XP_026687734.1 | $7 \times 10^{-70}$ |
| ERR-like 2 | XM_026822495 | XP_026678296 | $6 \times 10^{-113}$ |

### *Drosophila* display increased death and aberrant developmental timing when fed PT compounds

At a concentration of 1 mg mL^−1^, the PT150 and PT155 produced 100% overall lethality (60% in pupation and 40% as larvae). Compounds PT159 and PT160 demonstrated 100% lethality within 24 hours.

At a dosage of 0.1 mg mL^−1^, increased lethality was observed for all treated groups.

### Increased Scleratization Observed Following Exposure to PT Compounds

In the course of feeding PT155 to larvae to assess lethality, several pupae and larvae were observed to have darkened areas, beyond what was seen in control groups (representative images shown in Fig. 4). These dark areas are consistent with previously reported in several mutants of the ecdysone receptor (EcR) gene (Li & Bender, 2000). Although relatively few larvae showed this phenotype (3 out of 10), this phenotype is of interest, due to the previously reported interaction between EcR and GRE.

**Figure 1.**
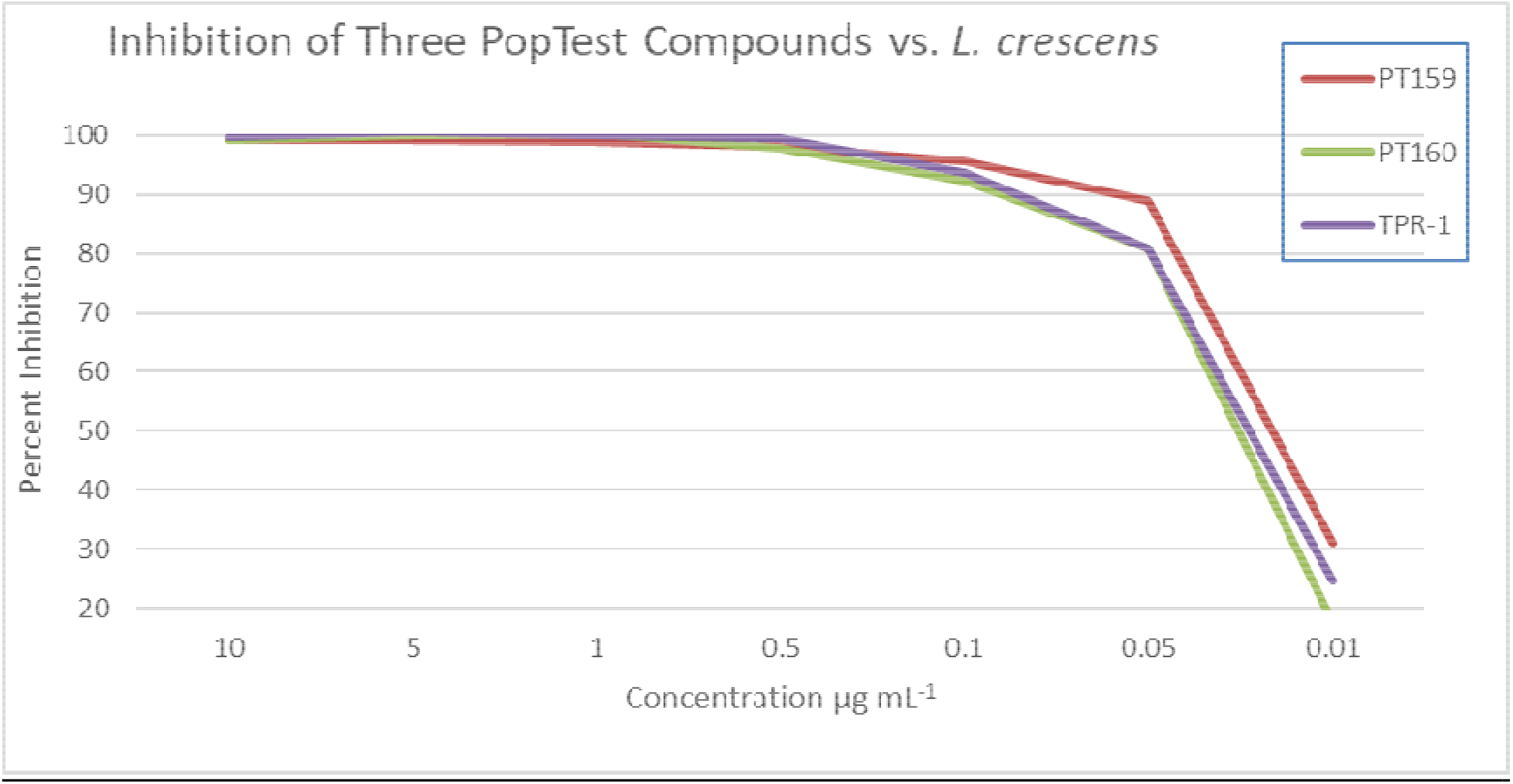
Effectiveness of PT Compounds Against *L. crescens*. PT compounds were assessed from 10 to 0.01μg mL^−1^ to assay their ability to inhibit bacterial growth.

**FIgure 2.**
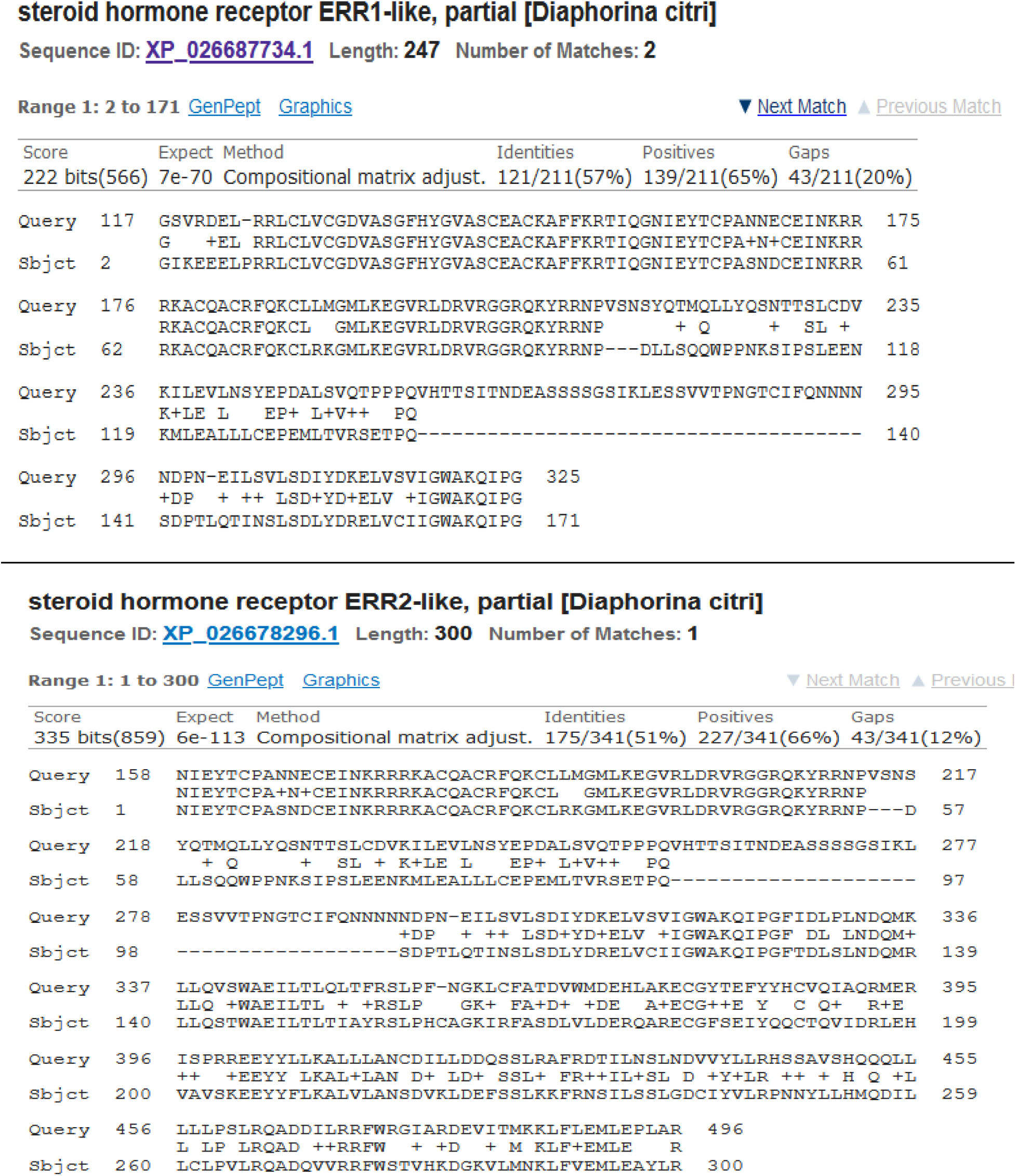
BLAST Alignments Between *D. citri* ERR-like Proteins and *Drosophila* ERR.

**Figure 3.**
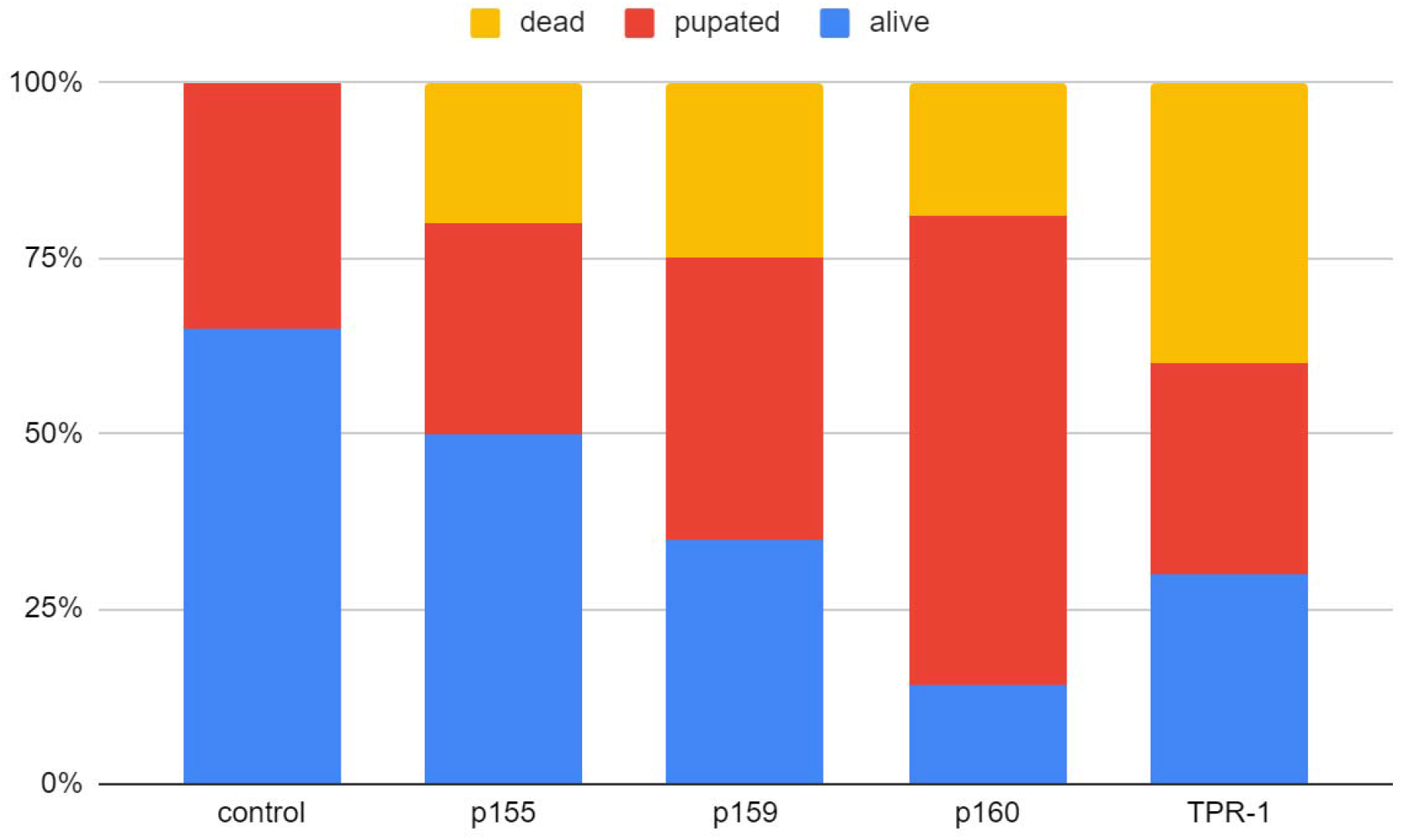
PT compounds increase lethality and induce aberrant pupation when compared to controls. 3rd instar foraging larvae were fed PT compounds at a concentration of 0.1mg mL^−1^, and their survival and development were assessed after 48 hours. All PT compounds increased the number of pupae present when compared to controls, and all displayed increased lethality.

**Figure 4.**
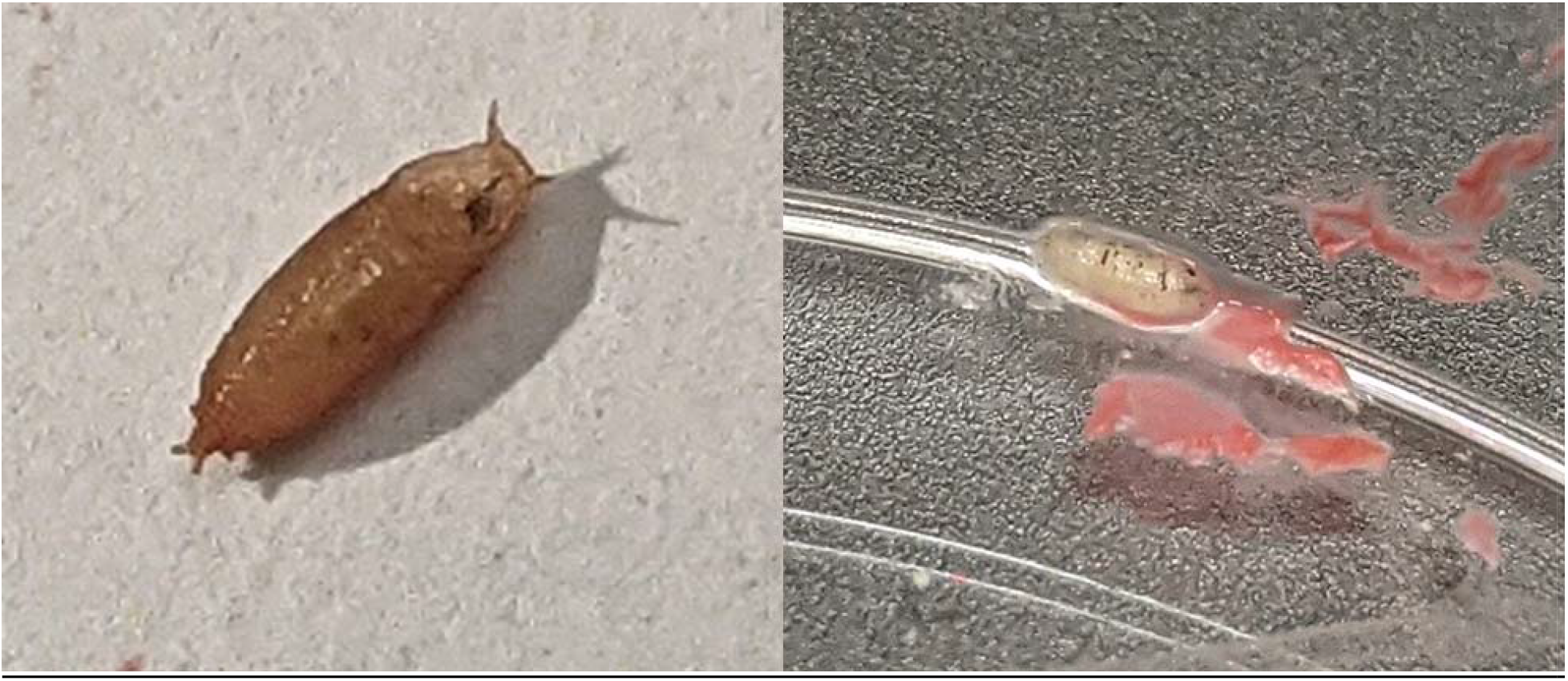
PT155 induced aberrant scleratization in a small number of treated individuals. Two representative images shown here, one of a pupa (left) with dark spots visible through the pupal casing, and one larva (right) shown with dark patches and stripes along segment boundaries, as well as sporadically between the segments.

### PT compounds elevate glucose levels in pupae compared to controls

Pupae from the dosage regimen of 0.1 mg mL^−1^ were assayed for glucose levels. These pupae had been fed PT compounds over the course of 48 hours, and were individually measured. All PT compounds induced elevated glucose levels relative to control pupae at the same developmental stage (Fig. 5)

**Figure 5.**
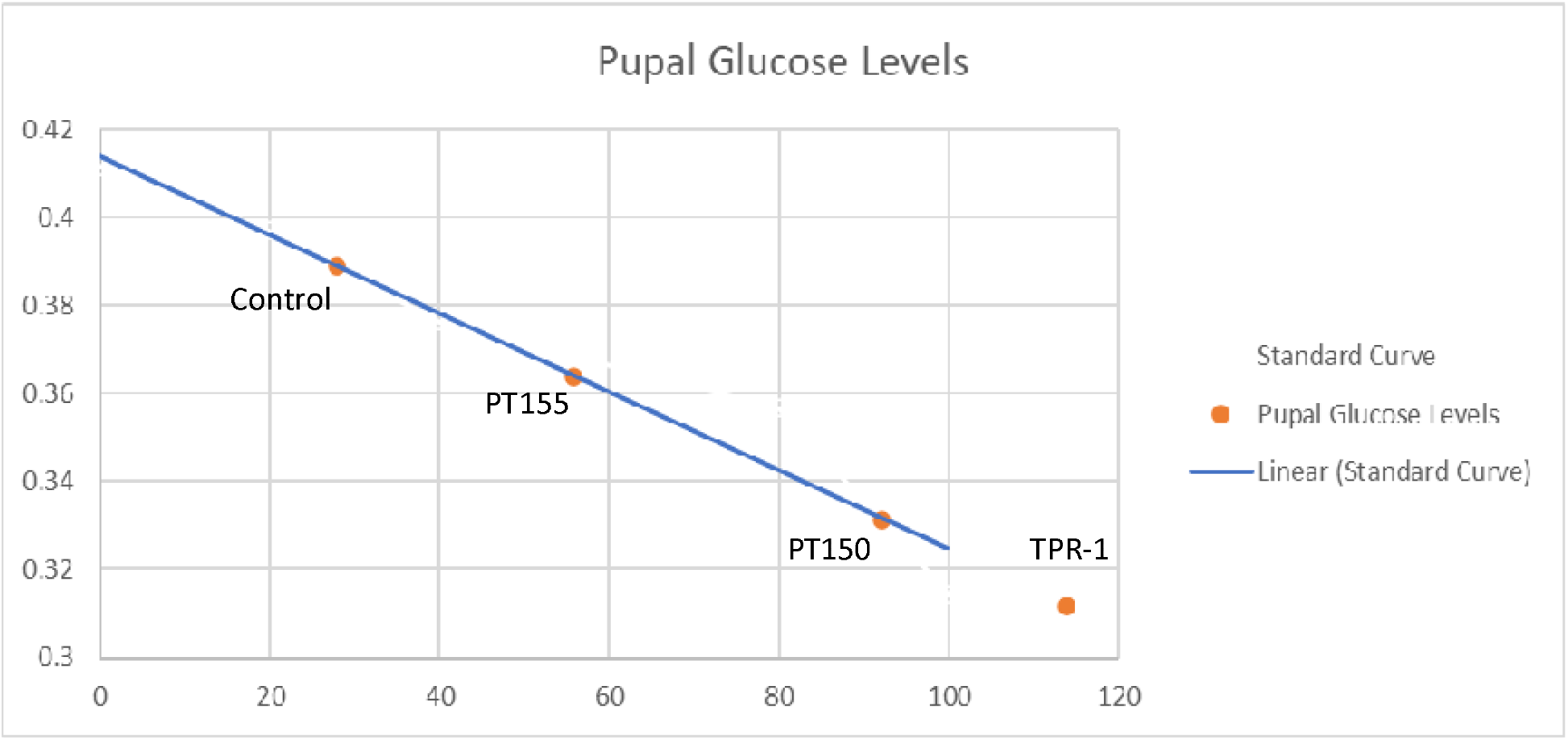
Results of glucose levels in larvae treated at 0.1 mg mL^−1^. Glucose levels were elevated in response to treatment; control larvae had an average value of 30 uM compared against treated larvae, PT155 (56 uM), PT150 (92 uM) and TPR-1 (114 uM). TPR-1 value was estimated from standard curve, as it was off-scale.

## Discussion

The damage to the Florida industry has become so severe that it has been estimated to become nonproductive in the next 10-15 years (Percy, 2017). In search of an answer, we offer evidence that a combination therapy of the rifampicin-derivative TPR-1 and glucocorticoid antagonists will sever the lifecycle of HLB simultaneously at the pathogen and transmission vector. The added temperature stability and light insensitivity of TPR-1 will enhance longevity and effectiveness in the field. Further, we anticipate that the effects of an antibiotic on the psyllid gut will make it inhospitable to transmit *Las*.

Antagonism of the ERR transcription factor in *Drosophila* was established by evidence of delayed larval development and increased lethality as well as by increased glucose level in treated versus untreated larvae. The genetic homologies between ERR and the ERR-like 1 and ERR-like 2 proteins of *D. citri* lead us to hypothesize that the effects will translate as active antagonists of their homologous proteins.

Taken together, we propose that some combination of the mechanisms depicted in Figure 6 will provide a dual action formulation that will both cure infected trees and prevent reinfection by controlling the populations of infected psyllids feeding directly on treated trees.

**Figure 6.**
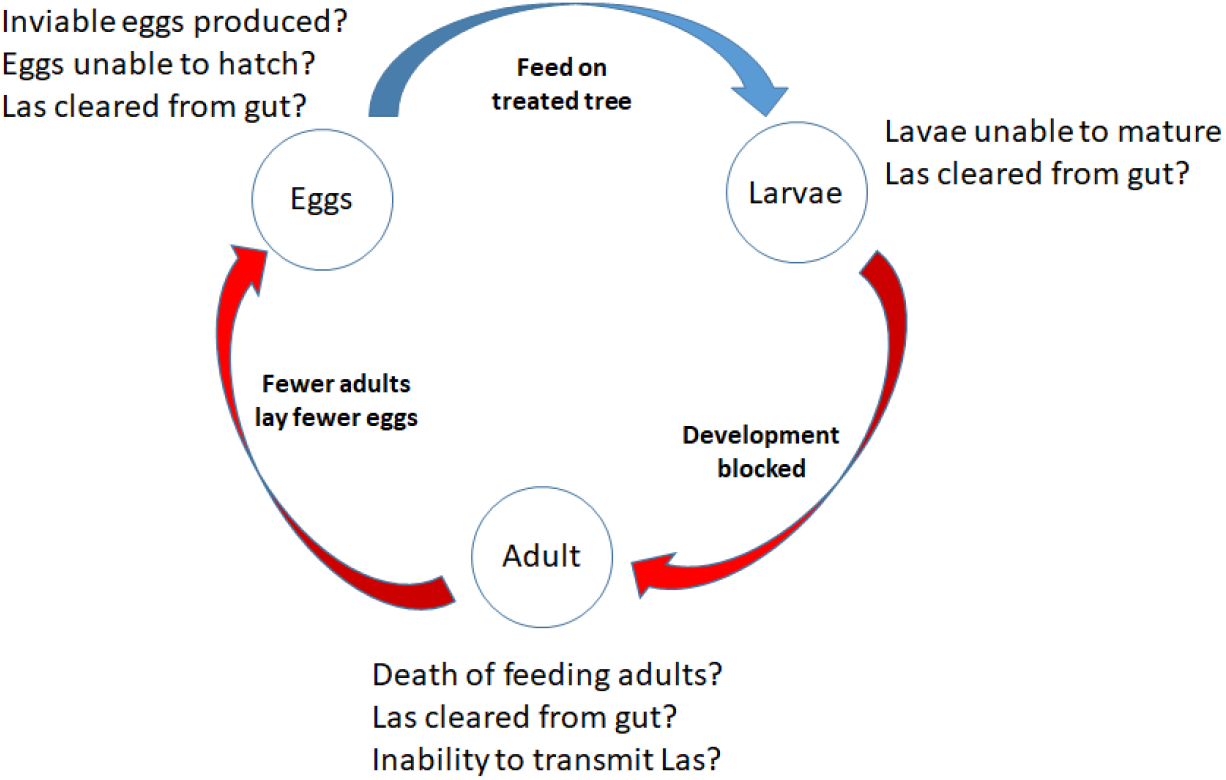
Model of how compounds will break the lifecycle of psyllids while curing infected trees

Much work remains to be done. Works in progress include phytotoxicity testing of compounds in citrus, establishment of an uninfected psyllid colony to verify mechanism translation and effects stemming from feeding on treated trees, establishment of effective compound levels in citrus and mechanisms for application. With the predicted final decade of Florida citrus looming as a consequence of inaction, it is our hope that this research will provide Florida citrus growers with the means to re-establish their livelihood and continue to supply citrus to the United States and the world. Globally, we hope our work will let other countries begin to recover from HLB damage as well. And finally, in areas, such as California, where HLB has not yet established a presence, we hope our research will provide the barrier that stops HLB.

## Abbreviations

*Las*: *Liberibacter asiaticus*
*Lc*: *Liberibacter crescens*
PT: Palisades Therapeutics
ERR: estrogen-related receptor
GREs: glucocorticoid response elements

## Acknowledgments

We would like to thank Palisade Therapeutics for providing their proprietary compounds tested in these studies.

